# Cell surface mechanics gate stem cell differentiation

**DOI:** 10.1101/798918

**Authors:** Martin Bergert, Sergio Lembo, Danica Milovanović, Mandy Börmel, Pierre Neveu, Alba Diz-Muñoz

**Affiliations:** Cell Biology and Biophysics Unit, European Molecular Biology Laboratory, Meyerhofstrasse 1, 69117 Heidelberg, Germany; Electron Microscopy Core Facility, European Molecular Biology Laboratory, Meyerhofstrasse 1, 69117 Heidelberg, Germany

**Author notes:** These authors contributed equally to this work (Martin Bergert, Sergio Lembo).

## Abstract

Cell differentiation typically occurs with concomitant shape transitions to enable specialized functions. To adopt a different shape, cells need to change the mechanical properties of their surface. However, whether conversely cell surface mechanics control the process of differentiation has been relatively unexplored. Here, we show that membrane mechanics gate the exit from naïve pluripotency of mouse embryonic stem cells. By measuring membrane tension during differentiation, we find that naïve stem cells release their plasma membrane from the underlying actin cortex when transitioning to a primed state. By mechanically tethering the plasma membrane to the cortex with a synthetic signalling-inert linker, we demonstrate that preventing this detachment forces stem cells to retain their naïve pluripotent state. We thus identify a decrease in membrane-to-cortex attachment as a new cell-intrinsic mechanism that is essential for stem cells to exit pluripotency.

---

During the development of most multicellular organisms, relatively round totipotent cells give rise to differentiated cells with all the dramatically different morphologies present in the adult body. Acquisition of fate and changes in cell shape often emerge concurrently. Cell shape is determined by surface mechanics and interactions with the extracellular environment. While cell-matrix interactions have been shown to be necessary (*1*, *2*) and in some cases sufficient (*3*) for differentiation, whether and how intrinsic surface mechanics regulate cell fate remains unknown. A particularly striking example of shape and identity change is the transition from the naïve to the primed pluripotent state in the widely used mouse embryonic stem cell (mESC) model. Naïve cells grow in compact colonies (**Fig. 1A-C**), and maintain this state when cultured in the presence of 2i/LIF (2 small molecule inhibitors [PD0325901 and CHIR99021] and leukemia inhibitory factor [LIF]). 2i/LIF removal provides a differentiation cue and the naïve colonies rapidly flatten into a monolayer of primed cells that grow lamellipodia-like protrusions in a process reminiscent of an epithelial-to-mesenchymal transition (*4*) (**Fig. 1A-C**, **Supplementary Movie 1**).

**Figure 1.**
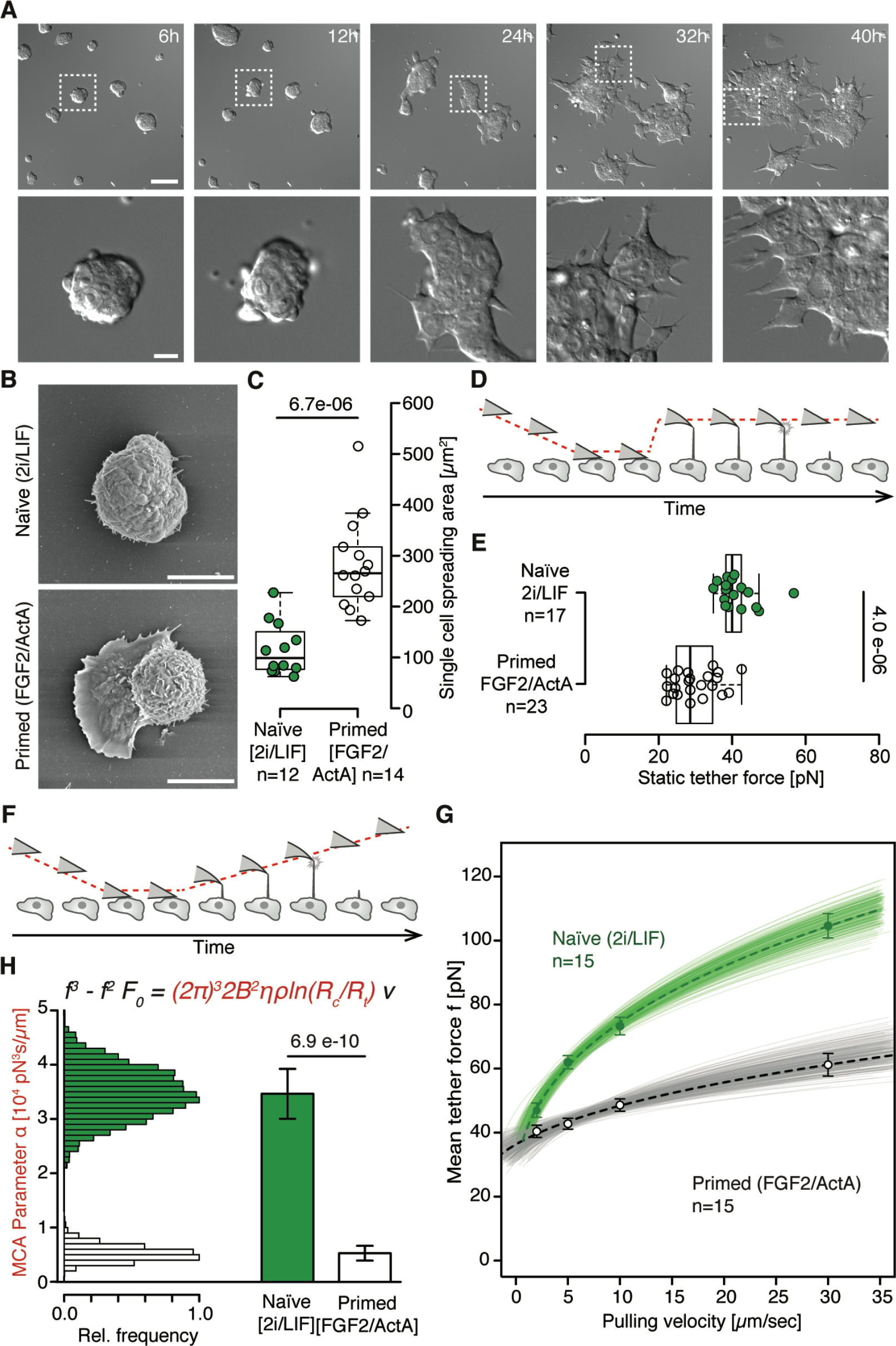
During exit from naïve pluripotency, cells spread with a concomitant reduction in apparent membrane tension and MCA. (**A**) Representative brightfield (DIC) images of mESC Rex1-GFPd2 cells during exit from pluripotency in N2B27 medium. Lower panel corresponds to a magnification of the boxed region in the upper panel. See also Supplementary Movie 1. Scale bar: 50 *μ*m (upper panel), 10 *μ*m (lower panel). (**B**) Representative scanning electron microscopy images of naïve (2i/LIF) and primed (FGF2/ActA) mESC Rex1-GFPd2 cells. Scale bar, 10 *μ*m. (**C**) Single cell spreading area quantified from scanning electron microscopy images. n: number of cells analysed. p-Value: Welch’s *t*-test. (**D**) Schematic of static tether pulling using atomic force spectroscopy. (**E**) Mean static tether force of naïve (2i/LIF) and primed (FGF2/ActA) mESC Rex1-GFPd2 cells. n: number of cells analysed in 2 independent experiments. p-Value: Mann-Whitney-U-Test. (**F**) Schematic of dynamic tether pulling using atomic force spectroscopy. (**G**) Force-velocity curve from dynamic tether pulling on naïve (2i/LIF) and primed (FGF2/ActA) mESC Rex1-GFPd2 cells. Data points are mean tether force f±SEM at 2, 5, 10 and 30 *μ*m/sec pulling velocity. Lines indicate Monte-Carlo fits of a model for dynamic tether formation (*8*) used to extract the MCA parameter *a* (see Materials and Methods for details). n: number of cells analysed in 3 independent experiments. (**H**) Monte-Carlo-based fitting of a model for dynamic tether formation (see equation, (*8*)) yields distributions for the MCA parameter *a*, used to determine mean and standard deviation of this parameter for each condition (see Materials and Methods for details). p-Value: *Z*-test.

Plasma membrane tension, or the energetic cost of stretching the plasma membrane, regulates several cellular processes, notably cell spreading (*5*) and migration, where it determines the size of the leading edge as well as the rate of lamellipodia extension (*6*, *7*). Given the striking protrusions stem cells form during exit to the primed state, we hypothesized that membrane tension may in fact play a role during the exit from naïve pluripotency. To test this, we measured apparent membrane tension by static tether pulling via single cell force spectroscopy, where a membrane tether is held by an AFM cantilever with a constant length until it breaks (**Fig. 1D**). Comparing naïve and primed cells, we found that the static tether force was significantly reduced in primed cells (from 41.3±5.25 to 30±5.92 pN, **Fig. 1E**). Such decrease in static tether force corresponds to an almost 50% reduction in apparent membrane tension (from 80 to 42 *μ*N/m; see Materials and Methods for details).

That primed cells have a lower membrane tension seems paradoxical given their shape (**Fig. 1B,C**), as leading edge growth and cell spreading are known to increase apparent membrane tension (*5*, *6*). Static tether pulling measures the combination of in-plane membrane tension (originating from the tight packing of hydrophobic lipid molecules to avoid contact with water molecules) as well as protein-mediated attachment to the underlying actomyosin cortex (termed membrane-to-cortex attachment or MCA), that also constrains membrane stretching ((*8*, *9*); reviewed in (*10*)). To determine which of these two mechanical parameters changes during stem cell differentiation, we measured specifically MCA by dynamic tether pulling, that measures the force required to extrude tethers across a range of different pulling speeds ((*8*, *11*); see Materials and Methods for details). We observed a dramatic 4-fold reduction of MCA during exit from naïve pluripotency (**Fig. 1F-H**). To exclude that this reduction was merely a consequence of cell spreading, we forced naïve cells to spread by plating them on Laminin 511 in the presence of 2i/LIF (**Supplementary Fig. 1A,B**) showing that an increase in cell area alone does not significantly affect MCA (**Supplementary Fig. 1C,D**). We can therefore conclude that cells strongly decrease the tethering of their plasma membrane to cortical actin when exiting pluripotency.

Given their co-occurrence, we next investigated whether the reduction in MCA was upstream, *i.e.* a regulator, or downstream, *i.e.* a consequence, of the exit from naïve pluripotency. To this end, we expressed <u>c</u>onstitutively <u>a</u>ctive Ezrin (CAEzrin, T567D (*12*)) in an inducible manner in naïve mESCs (**Supplementary Fig. 2A,B**). Ezrin links the plasma membrane to the underlying cortex and CAEzrin is the current gold standard in the field to experimentally increase MCA (*13*, *14*). Upon CAEzrin expression, mESC cells maintained a high MCA 48h after 2i/LIF removal (**Supplementary Fig. 2C, D**). To determine if expression of CAEzrin had affected the exit from naïve pluripotency, we assayed the presence of Rex1-GFPd2, a naïve marker which is down-regulated within 24h after 2i/LIF removal (*15*, *16*). Indeed, 48 h after 2i/LIF removal, CAEzrin expressing cells retained high levels of Rex1, very similar to naïve stem cells and in contrast to control cells expressing only mCherry (**Supplementary Fig. 2E**), suggesting that CAEzrin expression forces stem cells to retain a naïve pluripotent state.

In addition to mechanically linking the plasma membrane to the cortex, Ezrin also has biochemical roles in several signalling cascades (reviewed in (*17*)). To test if the striking effect on the exit from naïve pluripotency was truly a cell surface mechanical effect, we engineered a synthetic molecular tool that can directly link the plasma membrane to actin but is inert regarding signalling (iMC-linker, **Fig. 2A**). iMC-linker consists of a minimal actin binding domain (from Utrophin), fused to mCherry for fluorescence visualization and tagged with a lipidation consensus sequence (from Lyn) for plasma membrane insertion (for details see Materials and Methods). As predicted, iMC-linker, as well as its individual components, localized to the cell surface when expressed in mESCs in an inducible manner (**Fig. 2B, Supplementary Fig. 3A,B)**. Strikingly, inducing iMC-linker expression allowed stem cells to maintain a high MCA even 48h after triggering differentiation by 2i/LIF removal (**Fig. 2C**). Their MCA value measured by dynamic tether pulling was similar to naïve predecessor mESC and in stark contrast to the strongly decreased MCA of uninduced controls (**Fig. 2D**). To test if preventing the lowering of MCA with a synthetic mechanical linker was sufficient for cells to retain the naïve state, we again measured the down-regulation of the naïve marker Rex1-GFPd2 by flow cytometry. While stem cells expressing only the minimal actin binding or plasma membrane insertion motifs normally downregulated Rex1 during exit from naïve pluripotency, iMC-linker expressing cells maintained high expression of Rex1, suggesting that they remained naïve (**Fig. 3A,B**). To verify that our mechanical iMC-linker was truly able to keep stem cells in a naïve pluripotent state, we next performed a re-plating assay where primed cells do not robustly survive (**Fig. 3C**, for details see Materials and Methods). Indeed, iMC-linker expressing cells were able to generate 3-fold more colonies than controls expressing the two binding motifs separately (**Fig. 3D**). These results show that a synthetic mechanical tether between the plasma membrane and actin cortex is sufficient to force stem cells to retain a naïve pluripotent state, even two days after triggering differentiation. MCA is therefore a critical mechanical parameter that stem cells have to lower in order to progress from naïve to primed pluripotency. Together our findings uncover a fundamental cell-intrinsic mechanism whereby the regulation of cell surface mechanics controls cell identity.

**Figure 2.**
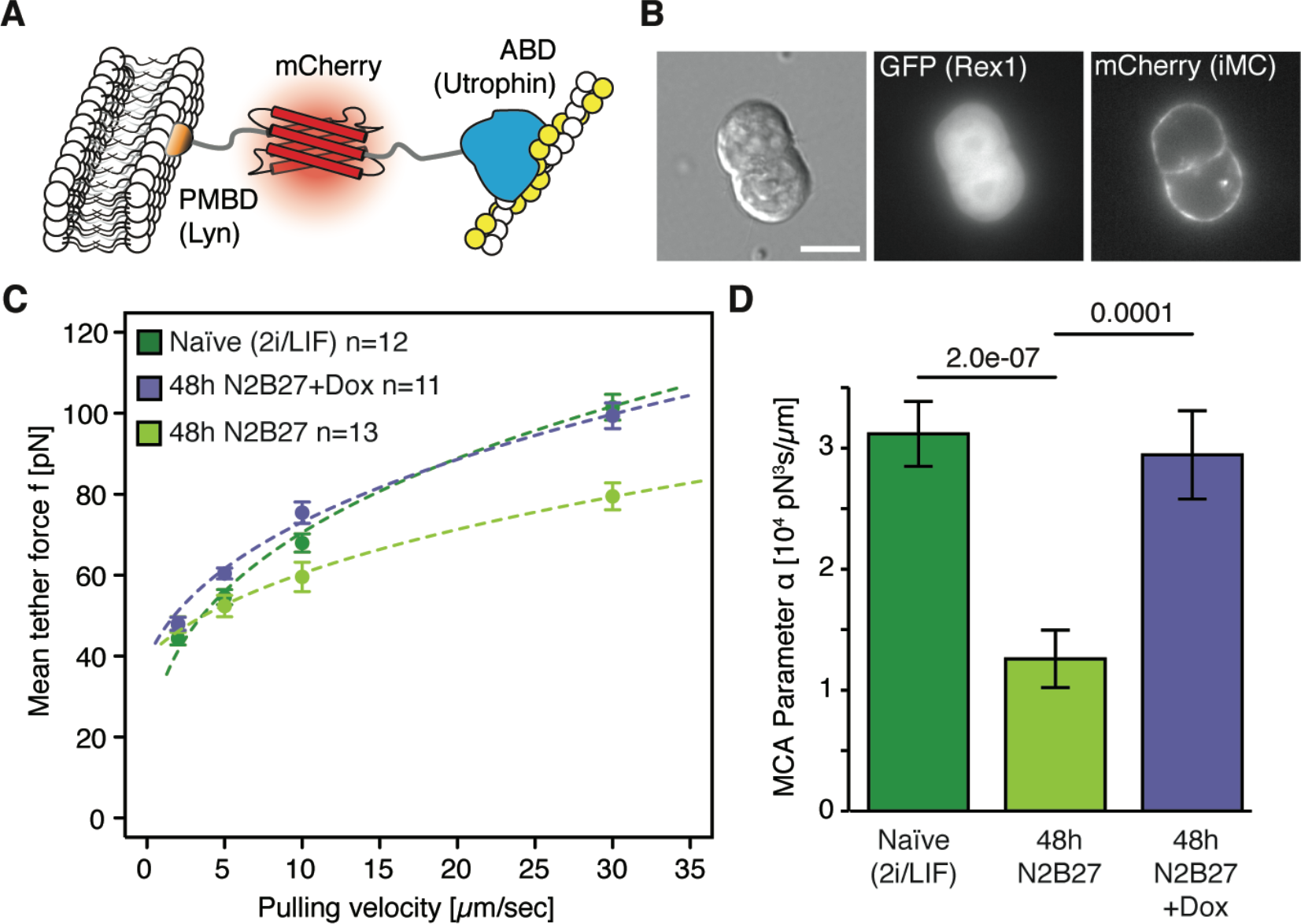
iMC-linker, an artificial signalling-inert MCA-protein, supresses the decrease of MCA during differentiation. (**A**) Schematic of the iMC-linker. (**B**) Representative brightfield (DIC) and fluorescent images of naïve mESC Rex1-GFPd2 *ind*-iMC cells expressing the iMC-linker in 2i/LIF+Dox medium. Scale bar: 10 *μ*m. (**C**) Force-velocity curve from dynamic tether pulling on mESC Rex1-GFPd2 *ind*-iMC cells in 2i/LIF medium and during exit from pluripotency in N2B27±Dox medium at 48 h. Data points are mean tether force f±SEM at 2, 5, 10 and 30 *μ*m/sec pulling velocity. n: number of cells analysed in 3 independent experiments. (**D**) Mean and standard deviation of the MCA parameter *a* obtained from Monte-Carlo-based fitting (see Materials and Methods for details). p-Value: *Z*-test.

**Figure 3.**
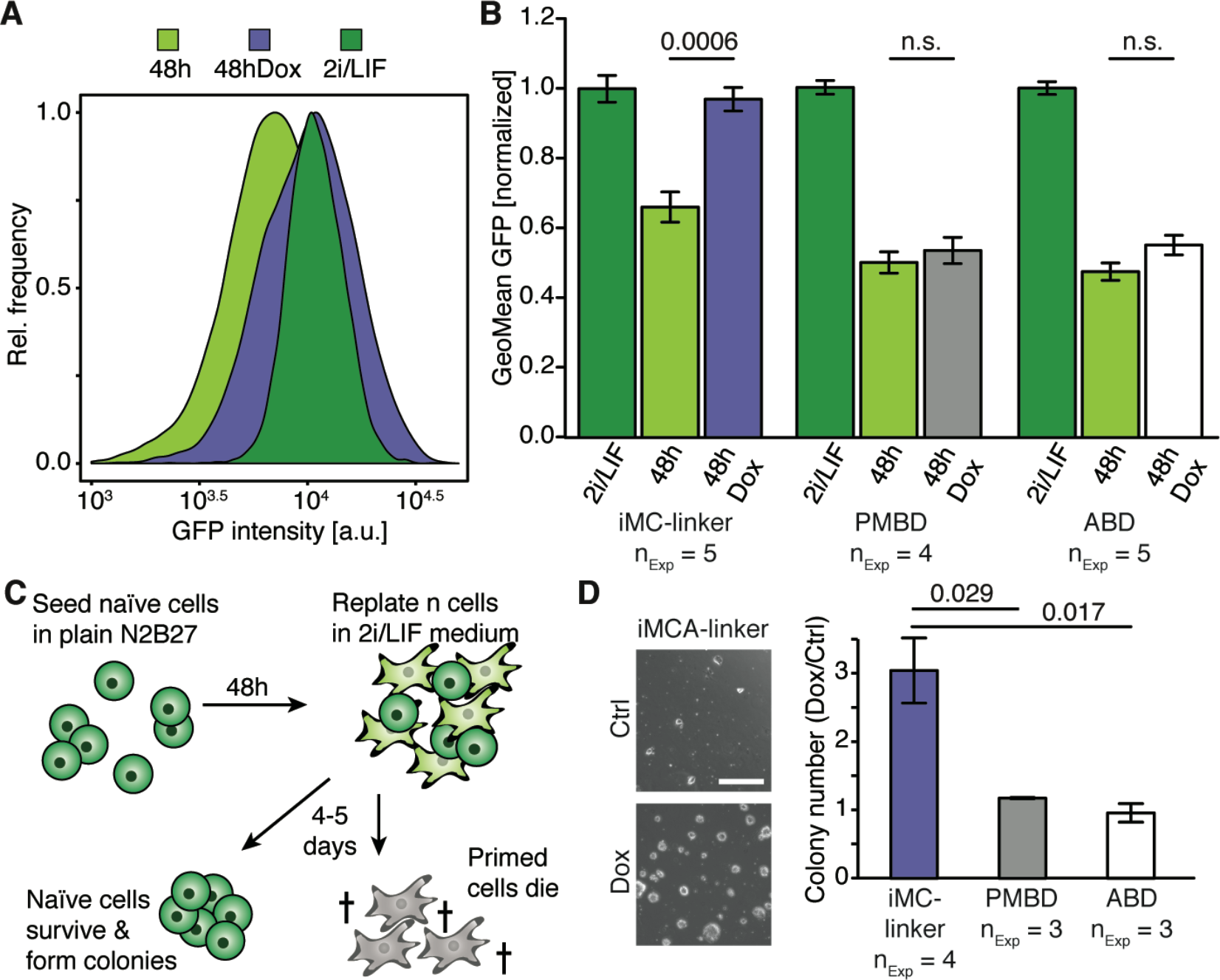
MCA gates the exit from naïve pluripotency. (**A**) Representative histograms of GFP intensity for mESC Rex1-GFPd2 *ind*-iMC cells in 2i/LIF medium and during exit from pluripotency in N2B27±Dox medium at 48 h. (**B**) Normalized GFP geometric mean intensities for mESC Rex1-GFPd2 *ind*-iMC, mESC Rex1-GFPd2 *ind*-PMBD and mESC Rex1-GFPd2 *ind*-ABD cells in 2i/LIF medium and during exit from pluripotency in N2B27±Dox medium at 48 h. n_Exp_: number of independent experiments. Error bars: SEM. p-Value: Welch’s *t*-test. (**C**) Schematic of the re-plating assay ((*37*, *38*), see Materials and Methods for details). (**D**) Left: Representative images of the re-plating assay for mESC Rex1-GFPd2 *ind*-iMC cells. Scale bar, 500 *μ*m. Right: Normalized colony number (Dox/Ctrl) for mESC Rex1-GFPd2 *ind*-iMC, mESC Rex1-GFPd2 *ind*-PMBD and mESC Rex1-GFPd2 *ind*-ABD cells re-plated after a 48 h exit in N2B27±Dox medium. n_Exp_: number of independent experiments. Error bars: SEM. p-Value: Welch’s *t*-test.

Changing cell surface mechanics by modifying the linking of the plasma membrane to the underlying actin cortex is likely to be important for embryonic stem cells also *in vivo*. Indeed, Ezrin, the classical endogenous protein that links the membrane to the cortex, marks the outside facing apical domain of 8-16 cell mouse embryos (*18*). This apical domain is necessary and sufficient for the first lineage segregation in early mouse embryos (*19*). Beyond the mouse, mechanical properties such as substrate stiffness, cortical contractility, fluid flow, cyclic stress, compression or luminal pressure have been linked to differentiation in several species (*3*, *20*–*30*). It will be interesting to explore in the future if MCA plays a regulatory role in those differentiation responses.

How exactly might lowering MCA gate the exit from naïve pluripotency? Previous studies have shown that membrane tension regulates several cellular functions such as endocytosis (*31*, *32*), phagocytosis (*33*), cell polarity (*6*) and the formation of blebs or lamellipodia (*11*, *34*). However, such studies have mostly used mechanical perturbations that also influence biochemical signalling and readouts of apparent membrane tension. Thus, the specific functions of cell surface signal transduction *versus* mechanics, as well as the relative contribution of the two membrane mechanics parameters, in-plane tension and MCA, have been impossible to disentangle. The synthetic iMC-linker we developed here as a precise tool to specifically manipulate MCA during stem cell differentiation should therefore prove useful to shed light on the mechanism of cell surface mechanics gated differentiation.

In conclusion, our work shows that changes of the mechanical properties of the cell surface, and in particular a lowering of MCA, are required for stem cells to exit naïve pluripotency. Moreover, together with reports of a role on lamellipodia initiation (*35*, *36*), it highlight MCA as a critical parameter for central cellular functions. Future studies, perhaps taking advantage of the novel iMC-linker tool we introduce here, will contribute to our understanding of how MCA controls cell signalling and cell shape during a wide range of cellular functions such as differentiation, polarity and migration.

## Acknowledgments

We thank Jan Ellenberg, Takashi Hiiragi, Chii Jou Chan and Erik Welf for critical reading of the manuscript. We are also grateful to Jamie Hackett for sharing plasmids, feedback along the project and critical reading of the manuscript. We thank the EMBL Flow cytometry core facility (and in particular Malte Paulsen) and the EMBL Electron microscopy core facility (and in particular Yannick Schwab) for assistance and advice.

## Funding

This study was funded by the European Molecular Biology Laboratory (EMBL) and the Human Frontiers Science Program (HFSP) grant number RGY0073/2018 (ADM); DM is the recipient of an Otto Bayer Fellowship from the Bayer Foundations. SL is a candidate for a joint Ph.D. between EMBL and Heidelberg University, Faculty of Biosciences.

## Author contributions

ADM, MBe and SL conceived the project and designed the experiments with advice from PN. MBe, SL, DM and MBö performed the experiments and analysed the data. ADM wrote the manuscript with input from SL, MBe and PN. All authors contributed to the interpretation of the data and read and approved the final manuscript.

## Supplementary information

### Supplementary Figures

**Supplementary Fig. 1.**
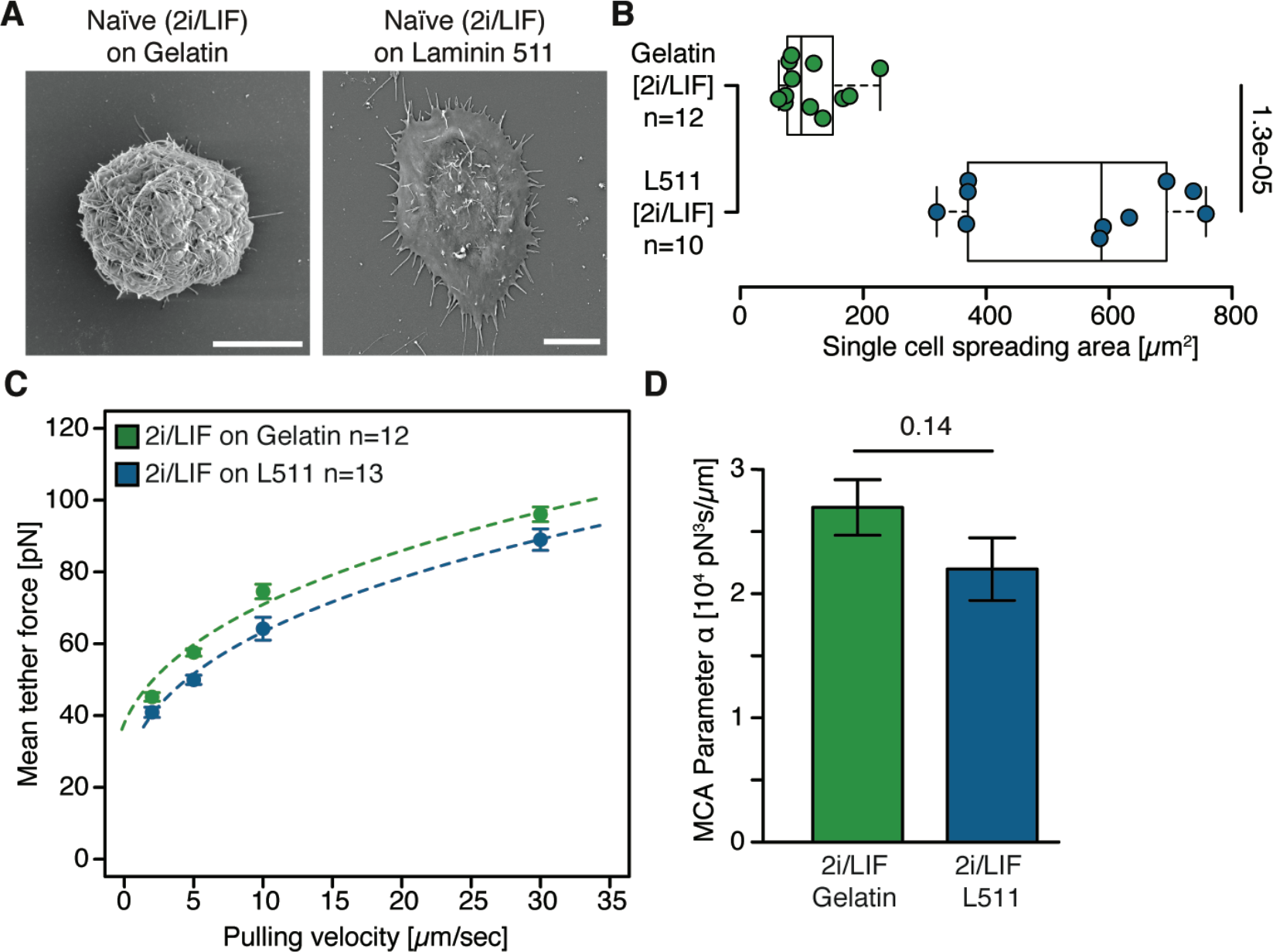
Cell spreading is not sufficient to decrease MCA. (**A**) Representative scanning electron microscopy images of naïve (2i/LIF) mESC Rex1-GFPd2 cells on Gelatin or on Laminin 511. Scale bar: 10 *μ*m. (**B**) Single cell spreading area quantified from scanning electron microscopy images. n: number of cells analysed. p-Value: Welch’s *t*-test. (**C**) Force-velocity curve from dynamic tether pulling on naïve (2i/LIF) mESC Rex1-GFPd2 cells plated on Gelatin or on Laminin 511. Data points are mean tether force f±SEM at 2, 5, 10 and 30 *μ*m/sec pulling velocity. n: number of cells analysed in 3 independent experiments. (**D**) Mean and standard deviation of the MCA parameter *a* obtained from Monte-Carlo-based fitting (see Materials and Methods for details). p-value: *Z*-test.

**Supplementary Fig. 2.**
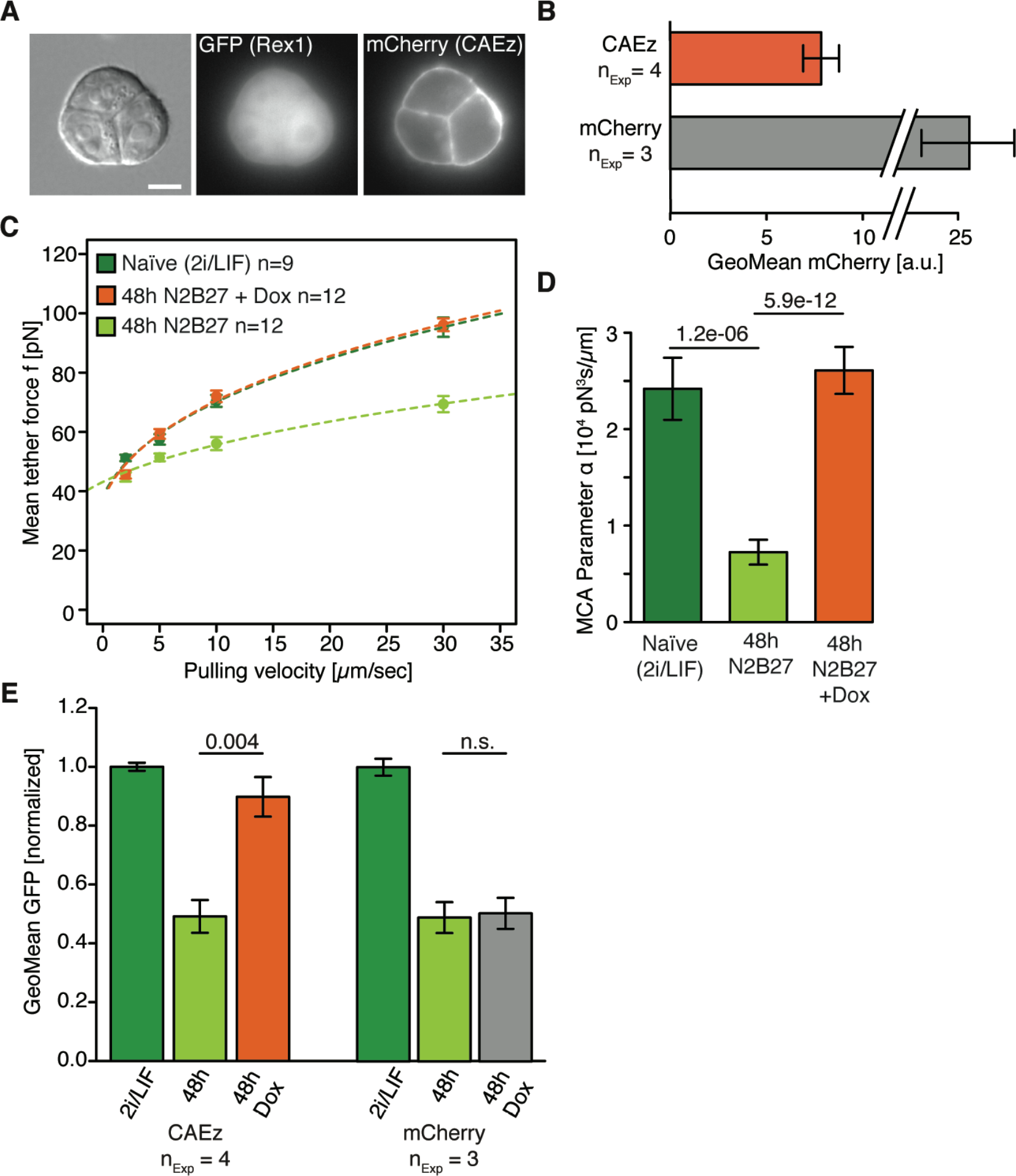
CAEzrin expression increases MCA and delays exit from naïve pluripotency. (**A**) Representative brightfield (DIC) and fluorescent images of naïve mESC Rex1-GFPd2 *ind*-CAEz cells expressing CAEz-mCherry in 2i/LIF+Dox medium. Scale bar: 10 *μ*m. (**B**) mCherry geometric mean intensities for mESC Rex1-GFPd2 *ind*-CAEz and mESC Rex1-GFPd2 *ind*-mCherry cells during exit from pluripotency in N2B27±Dox medium at 48 h. n_Exp_: number of independent experiments. Error bars: SEM. (**C**) Force-velocity curve from dynamic tether pulling on mESC Rex1-GFPd2 *ind*-CAEz cells in 2i/LIF medium and during exit from pluripotency in N2B27±Dox medium at 48 h. Data points are mean tether force f±SEM at 2, 5, 10 and 30 *μ*m/sec pulling velocity. n: number of cells analysed in 3 independent experiments. (**D**) Mean and standard deviation of the MCA parameter *a* obtained from Monte-Carlo-based fitting (see Materials and Methods for details). p-Value: *Z*-test. (**E**) Normalized GFP geometric mean intensities for mESC Rex1-GFPd2 *ind*-CAEz and mESC Rex1-GFPd2 *ind*-mCherry cells in 2i/LIF medium and during exit from pluripotency in N2B27±Dox medium at 48h. n_Exp_: number of independent experiments. Error bars: SEM. p-Value: Welch’s *t*-test.

**Supplementary Fig. 3.**
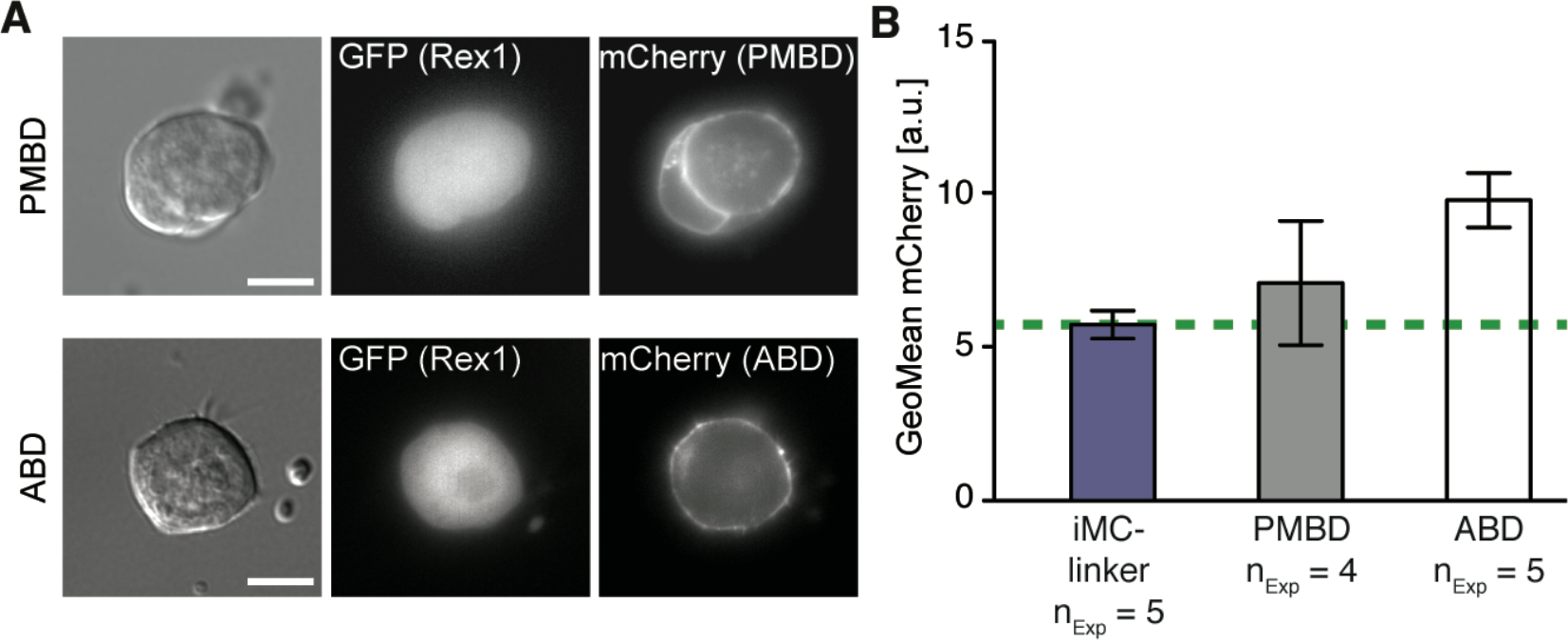
Localization of individual iMC-linker components and expression levels. (**A**) Representative brightfield (DIC) and fluorescent images of naïve mESC Rex1-GFPd2 *ind*-PMBD and mESC Rex1-GFPd2 *ind*-ABD cells in 2i/LIF+Dox medium. Scale bar: 10 *μ*m. (**B**) mCherry geometric mean intensities for mESC Rex1-GFPd2 *ind*-iMC, *ind*-PMBD and *ind*-ABD cells during exit from pluripotency in N2B27±Dox medium at 48 h. n_Exp_: number of independent experiments. Error bars: SEM.

### Supplementary Movies

**Supplementary Movie 1. Timelapse of the transition from naïve to primed pluripotency.** Scale bar: 50 μm. Time in hours:minutes after 2i/LIF removal.

## Materials and Methods

### Cell culture

mESC expressing Rex1-GFPd2 (*1*) were kindly provided by the Austin Smith’s laboratory (Cambridge Stem Cell Institute). Naïve cells were maintained in serum-free N2B27 medium containing 2i (1 *μ*M PD0325901and 3 *μ*M CHIR99021, both Tocris) and LIF (10 *μ*g/ml, EMBL Protein Expression Facility) (*2*) on polystyrene culture dishes coated with 0.1 % (w/v) Gelatin (Sigma) solution at 37 °C with 5 % CO_2_. N2B27 medium was prepared from a 1:1 mixture of DMEM/F12 (without HEPES, with L-glutamine) and neurobasal medium (no L-glutamine), supplemented with 0.5× B-27 (without vitamin A) and 0.5× N-2 supplement, 100 U/ml penicillin and 100 μg/ml streptomycin, 2.5 mM L-glutamine (all Thermofisher), 10 μg/ml BSA fraction V and 10 μg/ml human recombinant insulin (both Sigma). Medium was changed every other day and cells were passaged using 0.05 % Trypsin-EDTA (Thermofisher) at ratios of 1/4–1/10. To induce differentiation and the exit from pluripotency, cells were plated at a density of ~40.000/cm^2^ in plain N2B27 medium and cultured for 48 h unless otherwise stated.

### Re-plating assay

Cells were allowed to exit from pluripotency in plain N2B27 medium. After 48 h cells were resuspended and counted using trypan blue. A specific number of living cells (typical density: 5000-10.000/cm^2) was then re-plated in N2B27 2i/LIF. After 4-6 days, the number of colonies was manually counted. This assay quantifies the efficiency of pluripotency exit, as only naïve cells survive in 2i/LIF medium (*3*, *4*).

### Stable cell line generation

To generate stable cell lines with inducible constructs, sequences of interest were cloned into a Piggyback plasmid. Stable integration was achieved by co-transfecting the Piggyback plasmid and a plasmid encoding a transposase using Lipofectamine 3000 (Thermofisher), followed by selection with Geneticin. Single colony clones were finally screened for background expression levels and matching expression levels upon induction. To induce expression of the constructs, 1 μg/ml Doxycycline (Sigma) was added to the culture medium.

| Construct name | Construct description | Residues or mutations |
| --- | --- | --- |
| CAEz | constitutively active version of human Ezrin | T567D |
| <u>p</u>lasma <u>m</u>embrane-<u>b</u>inding <u>d</u>omain (PMBD) | human Tyrosine-protein kinase lyn lipidation motif | Residues 1-13 (5) |
| <u>a</u>ctin-<u>b</u>inding <u>d</u>omain (ABD) | Human utrophin actin binding domain (CH-CH) | Residues 1-246 (6) |
| iMC-linker | PMBD-mCherry-ABD | // |

### Flow cytometry

Cells were dissociated to single-cell suspension with 0.05 % Trypsin-EDTA (Thermofisher), resuspended in PBS supplemented with 0.1 % BSA and 2.5 mM EDTA, strained through a 40 *μ*m cell strainer (BD Biosciences) and analysed on an LSRFortessa flow cytometer (BD BioSciences). Flow cytometry data was gated on forward and side scatters using FlowJo software. Fluorescent levels of individual populations were quantified by their geometric means.

### Tether extrusion using atomic force spectroscopy

Apparent membrane tension and MCA were measured extruding plasma membrane tethers. In brief, OBL-10 cantilevers (Bruker) were mounted on a CellHesion 200 AFM (Bruker) which is integrated into an Eclipse Ti inverted light microscope (Nikon). Cantilevers were then calibrated using the thermal noise method (spring constant ~ 60 pN/nm) (reviewed in (*7*)) and coated with 2.5 mg/ml Concanavalin A (Sigma) for 1 h at 37 °C. Before the measurements, cantilevers were rinsed in PBS. For the measurements, cells were seeded on Cellview glass bottom dishes (Greiner) or 35 mm low *μ*-Dishes (Ibidi) filled with N2B27 medium with or without 2i/LIF according to the experiment.

Measurements were run as follows: approach velocity was set to 0.5 *μ*m/s while contact force and contact time were varied between 100 to 200 pN and 100 ms to 10 s respectively, aiming at maximizing the probability to extrude single tethers. Apparent membrane tension was measured using the static tether method: to ensure tether breakage at 0 velocity, the cantilever was retracted for 10 *μ*m at a velocity of 10 *μ*m/s. The position was then kept constant for 20 s and tether force at the moment of tether breakage was recorded at a sampling rate of 2000 Hz. MCA was measured using the dynamic tether method: each cell was probed multiple times at different retraction velocities (2, 5, 10, 30 *μ*m/s) in a random order; only tethers which broke during the retraction phase were considered. Tethers were allowed to retract completely between successive pulls. Resulting force–time curves were analysed using JPK Data Processing Software. Measurements were run at 37 °C and 5 % CO_2_ and samples were used no longer than 1 h for data acquisition.

### Tether data analysis and model assumptions

Static tether puling: apparent membrane tension (the sum of in-plane tension (*T*_*m*_) and MCA (*γ*)) depends on the breakage tether force (*f*_0_) and the bending rigidity (κ). We used a previously measured value for κ (2.7*10^−19^ Nm, (*8*)), which we assumed was unchanged upon exit from naïve pluripotency:

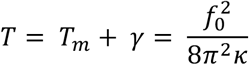

Dynamic tether pulling: To estimate the contribution of MCA, plasma membrane tethers were pulled at different retraction velocities (*v*), where the tether force (*f*) increases with increasing velocity. To interpret such measurements, the Brochard-Wyart *et al.* model was applied to the data using Monte-Carlo-based fitting (*9*):

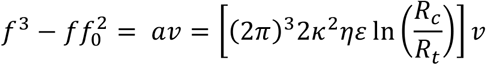

Since the radius of the cell (*R*_*c*_) ≫ radius of the tether (*R*_*t*_) and bending rigidity κ is assumed to be constant, the tether force increase with retraction velocity depends only on surface viscosity (η) and the density of the MCA linkers (ε). The term ηε thus reflects the effect of MCA on plasma membrane viscosity: the larger the extent of MCA, the higher is the drag of the lipids around integral membrane proteins connected to the underlying cortex. In this work, we report the parameter *a* as a proxy for MCA, as ηε is proportional to *a* and we assume *R*_*c*_, *R*_*t*_ and κ to be constant.

### Scanning electron microscopy

For scanning electron microscopy imaging, Rex1-GFPd2 mESC were cultured for 24 h on glass coverslips previously cleaned, plasma treated for 2 min and coated with 0.1 % Gelatin (Sigma), 10 *μ*g/ml human fibronectin (Corning) or 5 *μ*g/ml Laminin 511 (Biolamina). Cells were fixed for 30 min at room temperature with 4 % (w/v) formaldehyde (EMS) and 2.5 % (w/v) glutaraldehyde (EMS) dissolved in 0.1 M PHEM buffer (60 mM PIPES; 25 mM HEPES; 10 mM EGTA; 2 mM Magnesium chloride; pH 6.9). Afterwards, rinsing in PHEM buffer and water, post-fixation with 1 % (w/v) osmium tetroxide (EMS) in water and 0.8 % potassium hexacyanoferrate (III) (EMS) in water, 1 % (w/v) tannic acid (EMS) in water and 1 % (w/v) uranyl acetate (Serva) in water, dehydration in ascending series of ethanol and drying in ascending series of HMDS (Sigma-Aldrich) were performed using microwave-assisted processing (Biowave Pro, Pelco). Prior to imaging, a layer of gold was sputter-coated onto the sample (Quorum Q150RS). Cells were acquired with either the Teneo (Thermofisher) or the Crossbeam-540 (Zeiss) at an accelerating voltage of 5 kV detecting secondary electrons. Cell area was quantified manually using Fiji.

### Statistical analysis

Data were analysed, tested for statistical significance, fitted and visualized using R. No statistical method was used to predetermine sample size. No estimation of variance was performed. The Shapiro–Wilk test was used to test for normality of data. Welch’s *t*-test was chosen for statistical testing of normal distributed data with low sample size (n<30). For non-normal distributed data with low sample size (n<30), Mann–Whitney U-test was performed. For large sample sizes (n>30), the *Z*-Test was used.

## Bibliography

1. F. Chowdhury et al., Soft substrates promote homogeneous self-renewal of embryonic stem cells via downregulating cell-matrix tractions. PLoS ONE. 5, e15655 (2010).

2. P. Murray et al., The self-renewal of mouse embryonic stem cells is regulated by cell-substratum adhesion and cell spreading. Int J Biochem Cell Biol. 45, 2698–2705 (2013).

3. A. J. Engler, S. Sen, H. L. Sweeney, D. E. Discher, Matrix elasticity directs stem cell lineage specification. Cell. 126, 677–689 (2006).

4. C. Mulas et al., Defined conditions for propagation and manipulation of mouse embryonic stem cells (doi:10.1242/dev.173146). Development. 146, dev173146 (2019).

5. N. C. Gauthier, M.-A. Fardin, P. Roca-Cusachs, M. P. Sheetz, Temporary increase in plasma membrane tension coordinates the activation of exocytosis and contraction during cell spreading. Proceedings of the National Academy of Sciences. 108, 14467–14472 (2011).

6. A. R. Houk et al., Membrane tension maintains cell polarity by confining signals to the leading edge during neutrophil migration. Cell. 148, 175–188 (2012).

7. D. Raucher, M. P. Sheetz, Cell spreading and lamellipodial extension rate is regulated by membrane tension. J Cell Biol. 148, 127–136 (2000).

8. F. Brochard-Wyart, N. Borghi, D. Cuvelier, P. Nassoy, Hydrodynamic narrowing of tubes extruded from cells. Proc Natl Acad Sci USA. 103, 7660–7663 (2006).

9. F. M. Hochmuth, J. Y. Shao, J. Dai, M. P. Sheetz, Deformation and flow of membrane into tethers extracted from neuronal growth cones. Biophys J. 70, 358–369 (1996).

10. A. Diz-Muñoz, O. D. Weiner, D. A. Fletcher, In pursuit of the mechanics that shape cell surfaces. Nature Physics, 1–5 (2018).

11. A. Diz-Muñoz et al., Control of directed cell migration in vivo by membrane-to-cortex attachment. Plos Biol. 8, e1000544 (2010).

12. A. Gautreau, D. Louvard, M. Arpin, Morphogenic effects of ezrin require a phosphorylation-induced transition from oligomers to monomers at the plasma membrane. J Cell Biol. 150, 193–203 (2000).

13. Y. Liu et al., Constitutively active ezrin increases membrane tension, slows migration, and impedes endothelial transmigration of lymphocytes in vivo in mice. Blood. 119, 445–453 (2012).

14. C. Stefani et al., Ezrin enhances line tension along transcellular tunnel edges via NMIIa driven actomyosin cable formation. Nat Commun. 8, 1–14 (2017).

15. Y. Toyooka, D. Shimosato, K. Murakami, K. Takahashi, H. Niwa, Identification and characterization of subpopulations in undifferentiated ES cell culture. Development. 135, 909–918 (2008).

16. J. Wray et al., Inhibition of glycogen synthase kinase-3 alleviates Tcf3 repression of the pluripotency network and increases embryonic stem cell resistance to differentiation. Nat Cell Biol. 13, 838–845 (2011).

17. R. G. Fehon, A. I. McClatchey, A. Bretscher, Organizing the cell cortex: the role of ERM proteins. Nat Rev Mol Cell Biol. 11, 276–287 (2010).

18. S. Louvet, J. Aghion, A. Santa-Maria, P. Mangeat, B. Maro, Ezrin becomes restricted to outer cells following asymmetrical division in the preimplantation mouse embryo. Dev Biol. 177, 568–579 (1996).

19. E. Korotkevich et al., The Apical Domain Is Required and Sufficient for the First Lineage Segregation in the Mouse Embryo. Dev Cell. 40, 235–247.e7 (2017).

20. E. Farge, Mechanical induction of Twist in the Drosophila foregut/stomodeal primordium. Curr Biol. 13, 1365–1377 (2003).

21. J. Li et al., The Strength of Mechanical Forces Determines the Differentiation of Alveolar Epithelial Cells. Dev Cell. 44, 297–312.e5 (2018).

22. D. M. Cohen, C. S. Chen, Mechanical control of stem cell differentiation (2008), doi:10.3824/stembook.1.26.1.

23. A. Aguilar et al., Importance of environmental stiffness for megakaryocyte differentiation and proplatelet formation. Blood. 128, 2022–2032 (2016).

24. M. M. Pathak et al., Stretch-activated ion channel Piezo1 directs lineage choice in human neural stem cells. Proceedings of the National Academy of Sciences. 111, 16148–16153 (2014).

25. L. Przybyla, J. N. Lakins, V. M. Weaver, Tissue Mechanics Orchestrate Wnt-Dependent Human Embryonic Stem Cell Differentiation. Cell Stem Cell. 19, 462–475 (2016).

26. T. E. North et al., Hematopoietic stem cell development is dependent on blood flow. Cell. 137, 736–748 (2009).

27. J. R. Hove et al., Intracardiac fluid forces are an essential epigenetic factor for embryonic cardiogenesis. Nature. 421, 172–177 (2003).

28. C. J. Chan et al., Hydraulic control of mammalian embryo size and cell fate. Nature. 571, 112–116 (2019).

29. J.-L. Maître et al., Asymmetric division of contractile domains couples cell positioning and fate specification. 536, 344–348 (2016).

30. F. Chowdhury et al., Material properties of the cell dictate stress-induced spreading and differentiation in embryonic stem cells. Nature Materials. 9, 82–88 (2009).

31. N. C. Gauthier, O. M. Rossier, A. Mathur, J. C. Hone, M. P. Sheetz, Plasma membrane area increases with spread area by exocytosis of a GPI-anchored protein compartment. Mol Biol Cell. 20, 3261–3272 (2009).

32. C. Rauch, A.-C. Brunet, J. Deleule, E. Farge, C2C12 myoblast/osteoblast transdifferentiation steps enhanced by epigenetic inhibition of BMP2 endocytosis. Am J Physiol, Cell Physiol. 283, C235–43 (2002).

33. T. A. Masters, B. Pontes, V. Viasnoff, Y. Li, N. C. Gauthier, Plasma membrane tension orchestrates membrane trafficking, cytoskeletal remodeling, and biochemical signaling during phagocytosis. Proceedings of the National Academy of Sciences. 110, 11875–11880 (2013).

34. A. Diz-Muñoz et al., Steering cell migration by alternating blebs and actin-rich protrusions. BMC Biology. 14, 74–13 (2016).

35. E. S. Welf et al., A unified role for membrane-cortex detachment during cell protrusion initiation. BioRxiv. 1, 37–34 (2019).

36. A. Bisaria, A. Hayer, D. Garbett, D. Cohen, T. Meyer, Membrane proximal F-actin restricts local membrane protrusions and directs cell migration. BioRxiv. 50, 064002–21 (2019).

37. D. Cirera-Salinas, C. Ciaudo, Exit from Pluripotency Assay of Mouse Embryonic Stem Cells. BIO-PROTOCOL. 7, 21900–21905 (2017).

38. J. Betschinger et al., Exit from pluripotency is gated by intracellular redistribution of the bHLH transcription factor Tfe3. Cell. 153, 335–347 (2013).

## Supplementary Bibliography

1. J. Wray et al., Inhibition of glycogen synthase kinase-3 alleviates Tcf3 repression of the pluripotency network and increases embryonic stem cell resistance to differentiation. Nat Cell Biol. 13, 838–845 (2011).

2. C. Mulas et al., Defined conditions for propagation and manipulation of mouse embryonic stem cells (doi:10.1242/dev.173146). Development. 146, dev173146 (2019).

3. D. Cirera-Salinas, C. Ciaudo, Exit from Pluripotency Assay of Mouse Embryonic Stem Cells. BIO-PROTOCOL. 7, 21900–21905 (2017).

4. J. Betschinger et al., Exit from pluripotency is gated by intracellular redistribution of the bHLH transcription factor Tfe3. Cell. 153, 335–347 (2013).

5. D. A. Zacharias, J. D. Violin, A. C. Newton, R. Y. Tsien, Partitioning of lipid-modified monomeric GFPs into membrane microdomains of live cells. Science (New York, NY). 296, 913–916 (2002).

6. S. J. Winder et al., Utrophin actin binding domain: analysis of actin binding and cellular targeting. J Cell Sci. 108 (Pt 1), 63–71 (1995).

7. J. L. Hutter, J. Bechhoefer, Calibration of atomic-force microscope tips. Rev. Sci. Instrum. 64, 1868–1873 (1993).

8. F. M. Hochmuth, J. Y. Shao, J. Dai, M. P. Sheetz, Deformation and flow of membrane into tethers extracted from neuronal growth cones. Biophys J. 70, 358–369 (1996).

9. F. Brochard-Wyart, N. Borghi, D. Cuvelier, P. Nassoy, Hydrodynamic narrowing of tubes extruded from cells. Proc Natl Acad Sci USA. 103, 7660–7663 (2006).

